## Supplementary material for "Predicting the Structural Impact of Human Alternative Splicing": SuppFigs

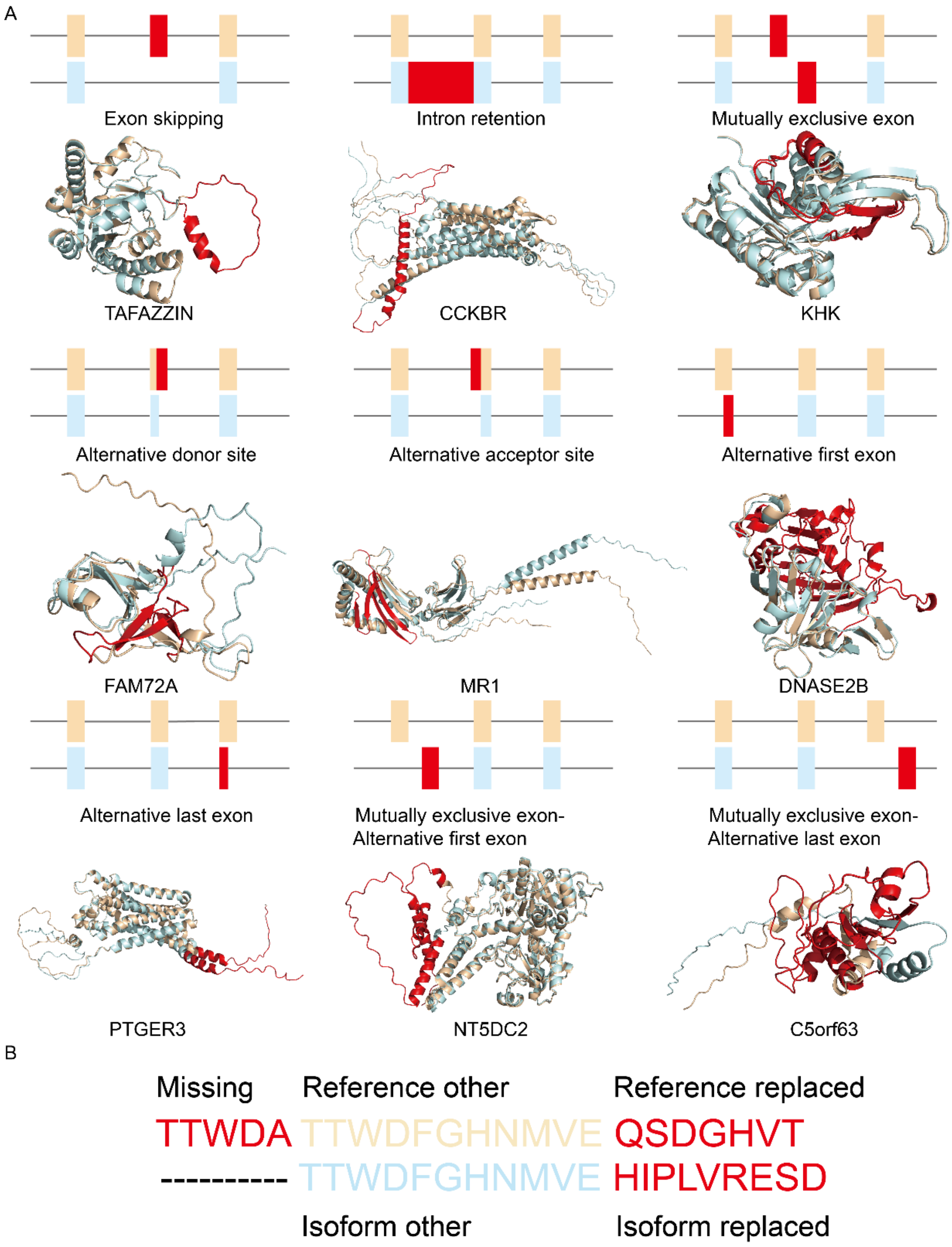


**Figure S1: Illustration of nine alternative splicing types and different alternative splicing regions.** (**A**) Diagram for nine alternative splicing types represented by transcripts and example structures. For each type, reference and isoform structures are colored in wheat and pale cyan, respectively. Alternatively spliced regions are shown in red. (**B**) Explanation of spliced and unspliced regions. Splicing regions including ‘Missing’, ‘Reference replaced’ and ‘Isoform replaced’ are colored in red, and unspliced regions (‘Reference other’ and ‘Isoform other’) are presented in wheat and pale cyan, respectively.

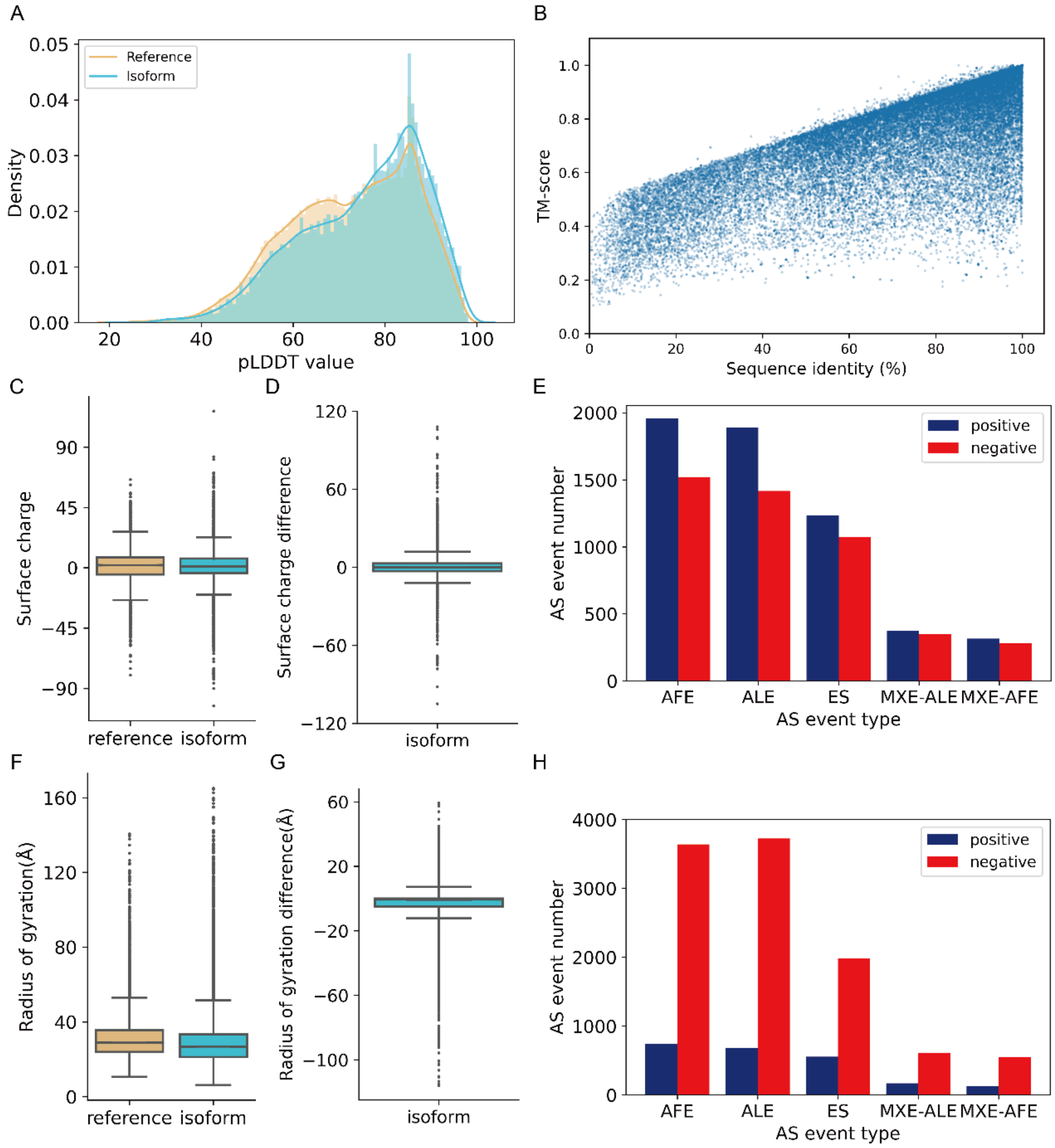


**Figure S2: Structural metrics analysis for the CHESS dataset.** (**A**) Distribution of prediction quality for 15,727 reference and 9,7824 isoform structures from the CHESS dataset. (**B**) Scatter plot of percent sequence identity between reference and alternate isoform (x-axis) vs. TM score (y-axis) for the CHESS dataset. (PCC=0.783). (**C**) Overall surface charge distribution between reference and isoform structures from the CHESS dataset (p-value: 6.053e-11 Mann–Whitney U test). (**D**) Differences of surface charge for the CHESS dataset. (**E**) The five most frequent alternative splicing events in the positive and negative surface charge outliers from the CHESS dataset. (**F**) Overall radius of gyration distribution between reference and isoform structures from the CHESS dataset (p-value: 1.074e-115 Mann–Whitney U test). (**G**) Differences of radius of gyration from the CHESS dataset. (**H**) The five most frequent alternative splicing events in the positive and negative radius of gyration outliers from the CHESS dataset.

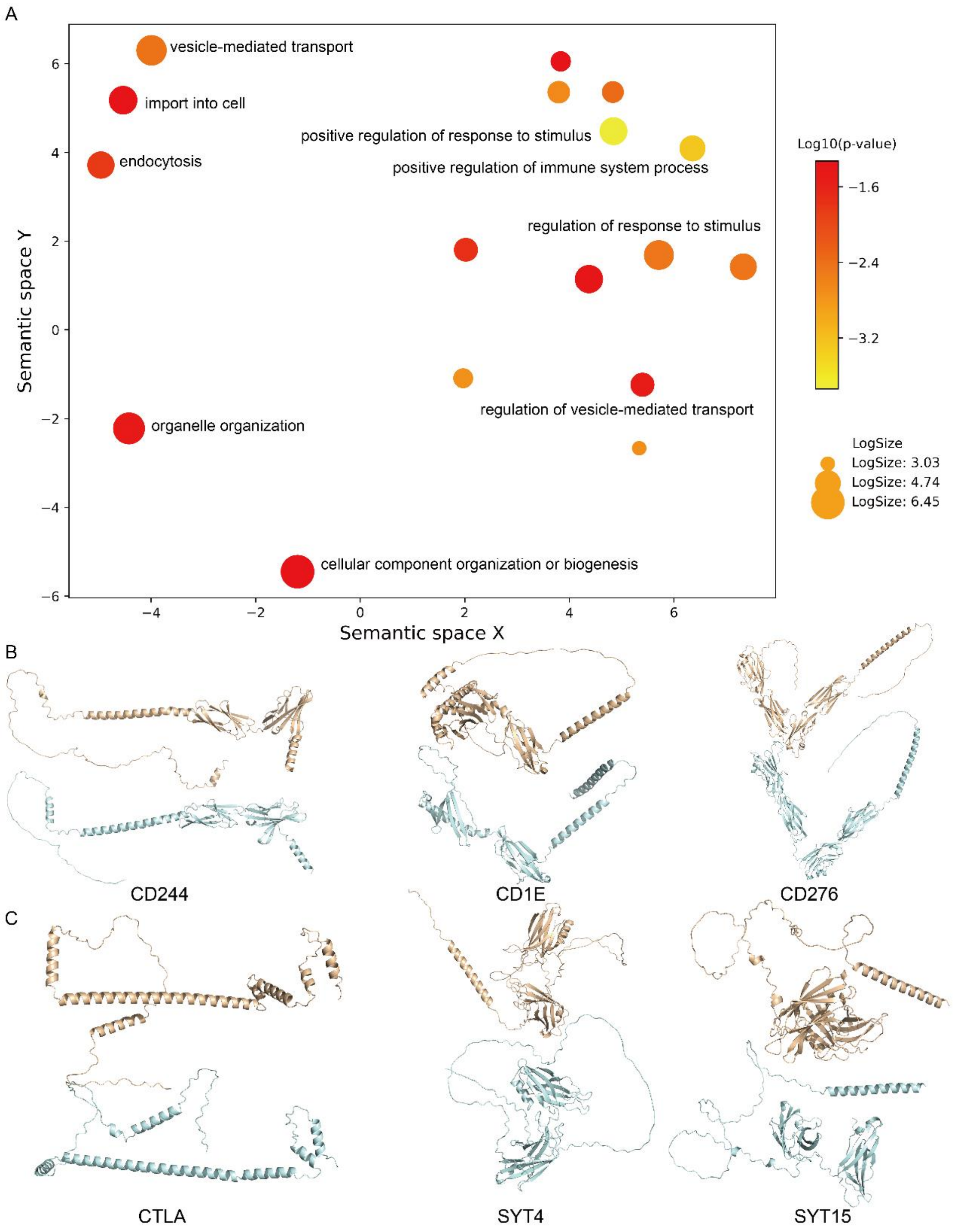


**Figure S3: Loop-affected high sequence identity but low TM-score examples.** (**A**) Gene ontology analysis for loop-affected splicing isoforms, main GO terms are labeled, colored by the log10 p-value from the gene enrichment test. (**B**) Structures for immune response related high identity but low TM-score examples: CD244, CD1E and CD276. (**C**) Structures for protein transport related high identity but low TM-score examples: CTLA, SYT4 and SYT15.

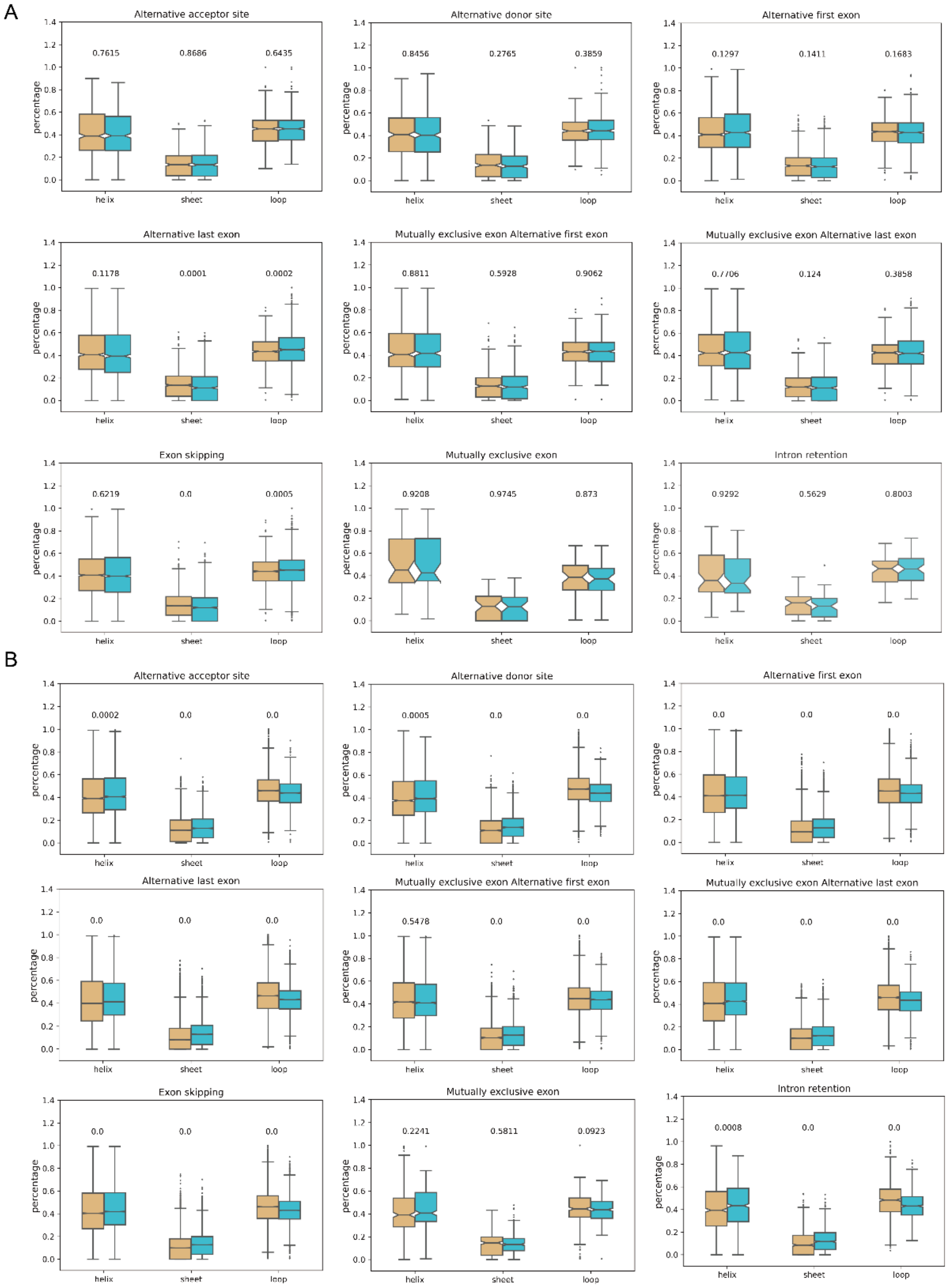


**Figure S4: Secondary structure percentage for Swissprot dataset (A) and CHESS dataset (B).** P-value labeled for each secondary structure percentage (helix, sheet and loop) between isoform and reference based on Mann–Whitney U test.

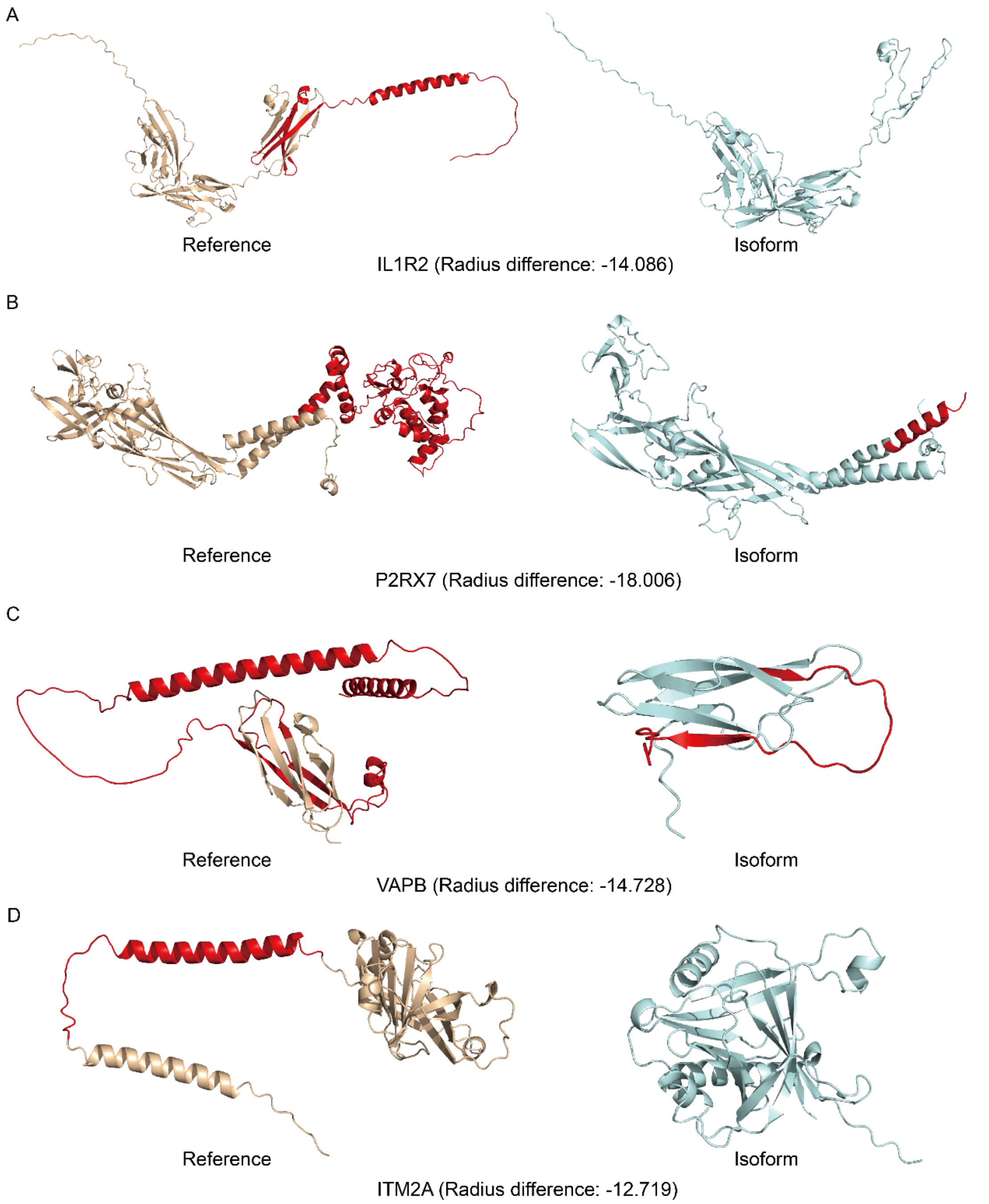


**Figure S5: Examples of negative radius of gyration outliers.** Reference and isoform structures for IL1R2 (A), P2RX7 (B), VAPB (C) and ITM2A (D) are presented in wheat and pale cyan. Alternatively spliced portions of the structure are colored red.

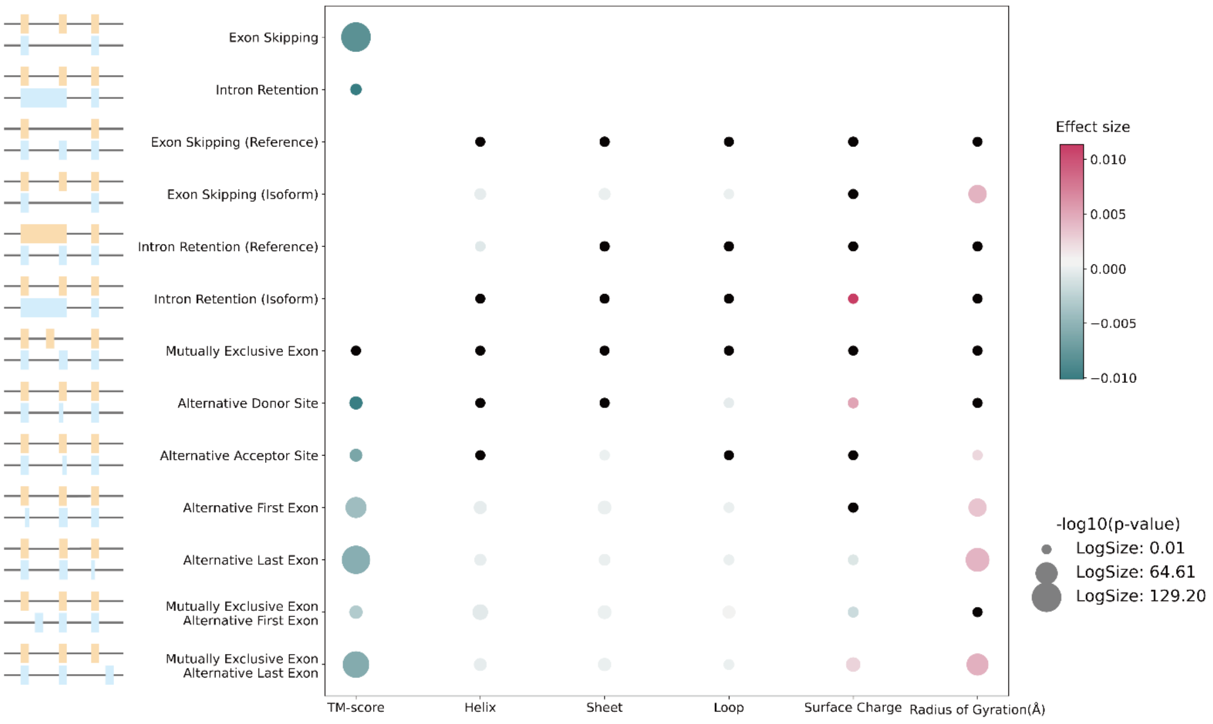


**Figure S6: Per-residue effect caused by nine alternative splicing types in terms of different structural metrics.** Effect size is colored from green to pink to represent the negative to positive per-residue effect. Dot size represents the negative log10 p-value, and dots with insignificant p-value (p-value>0.05) are colored in black. P-value for each effect size is computed from the lm function in R.

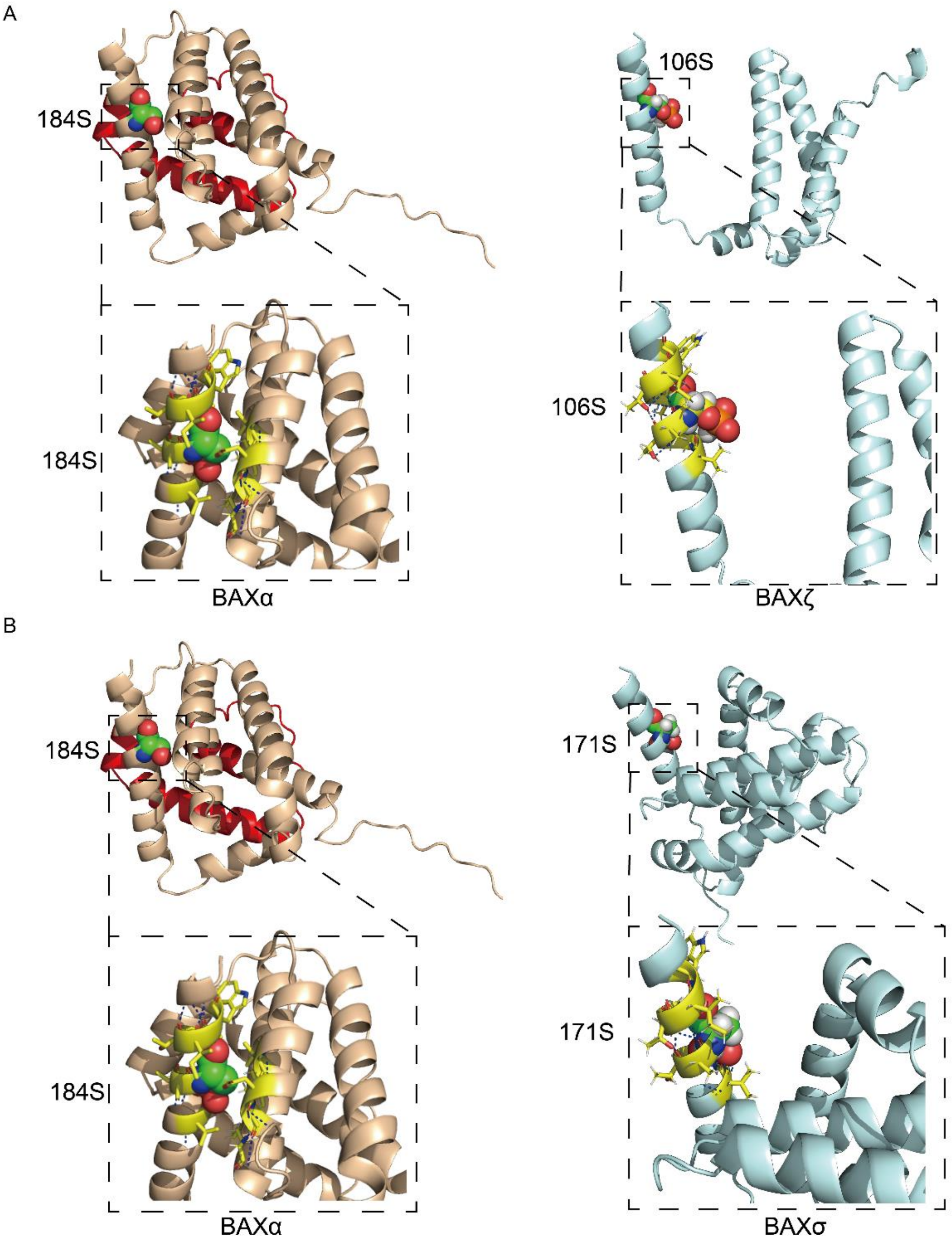


**Figure S7: Surrounding environment change of post translational modification (PTM) sites of BAXζ and BAXσ.** (**A**) Residue 184S is buried in BAXα (RSA: 1.18Å$^2$), and the corresponding PTM site 106S is exposed in BAXζ (RSA: 51.55Å$^2$). The residues within 4Å of the PTM site are represented in yellow sticks, and the polar contacts with those residues are represented in blue lines. (**B**) Residue 184S is buried in BAXα (RSA: 1.18Å$^2$), and the corresponding PTM site 171S is exposed in BAXσ (RSA: 53.52Å$^2$).

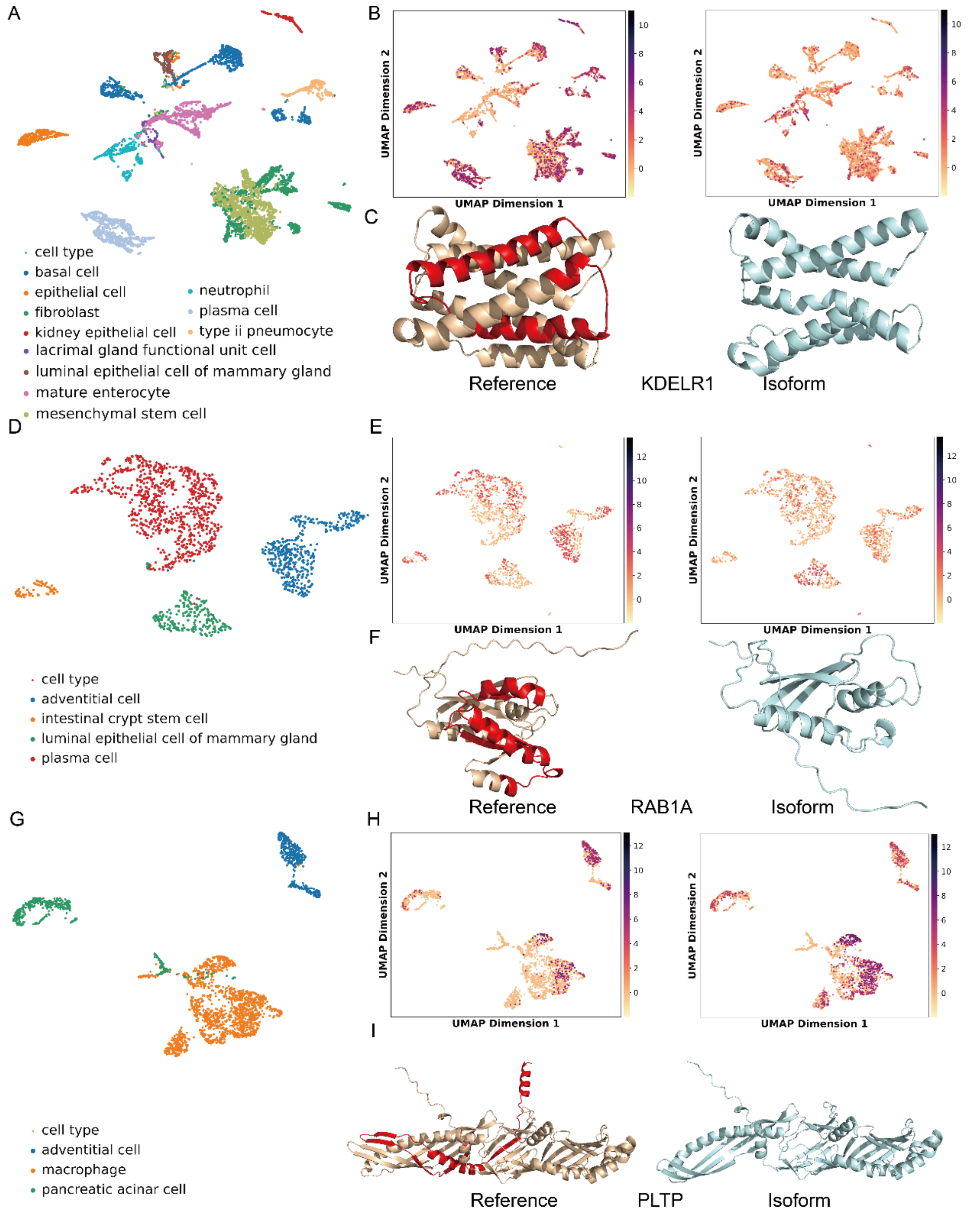


**Figure S8: Isoform usage shift examples discovered from scRNA-seq data.** Cell type clusters with isoform usage shift for KDELR1 (**A**), RAB1A (**D**) and PLTP (**G**) represented by UMAP plot. Expression of reference (left) and isoform (right) for KDELR1 (**B**), RAB1A **(E**) and PLTP (**H**). Reference and isoform structures for KDELR1 (**C**), RAB1A (**F**) and PLTP (**I**).

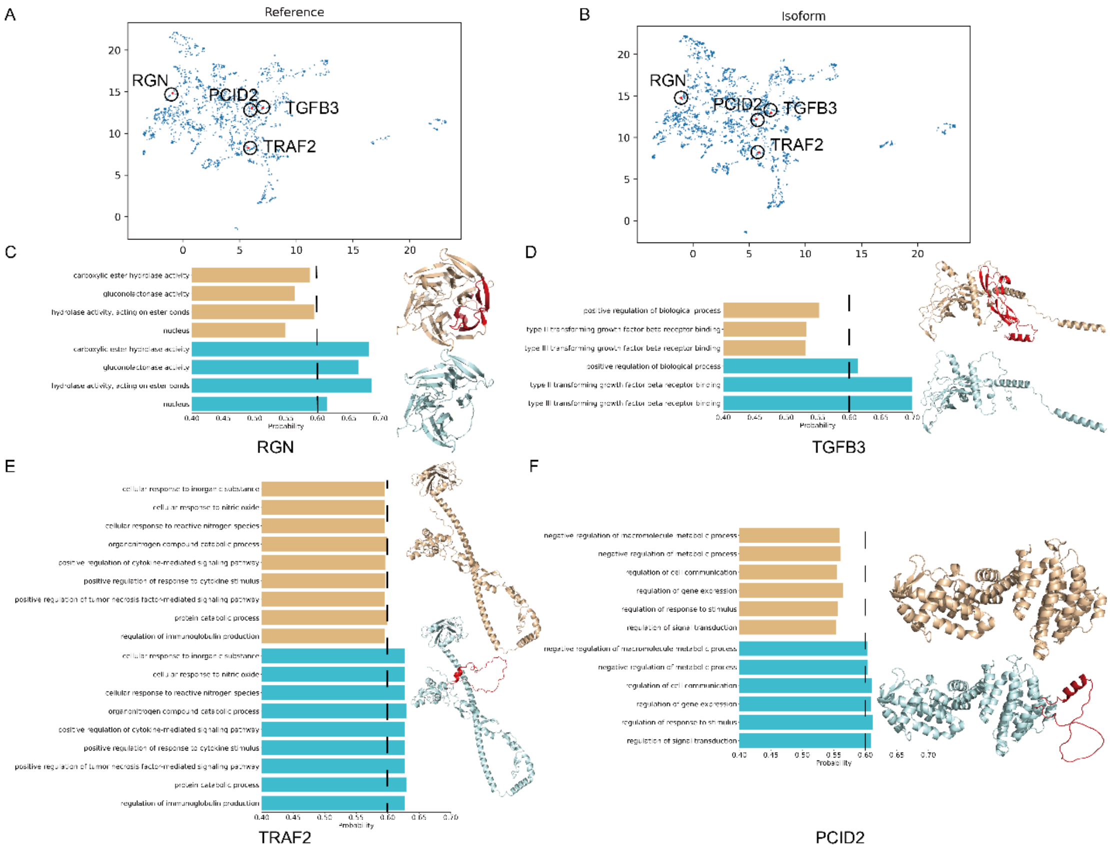


**Figure S9: Examples of gain of function spliced isoforms.** UMAP plot based on jaccard distance for reference (**A**) and isoform (**B**), and the gain of function examples are circled. Structures and the predicted gain-in go terms for RGN (**C**), TGFB3 (**D**), TRAF2 (**E**) and PCID2 (**F**), the alternative splicing regions are colored in red in the structures.
